# Enzymatic reactions dictated by the 2D membrane environment

**DOI:** 10.1101/2024.08.22.609272

**Authors:** Ru-Hsuan Bai, Chun-Wei Lin

## Abstract

The cell membrane is a fundamental component of cellular architecture. Beyond serving as a physical barrier that encloses the cytosol, it also provides a crucial platform for numerous biochemical reactions. Due to the unique two-dimensional and fluidic environment of the membrane, reactions that occur on its surface are subject to specific physical constraints. However, the advantages and disadvantages of membrane-mediated reactions have yet to be thoroughly explored. In this study, we reconstitute a classic proteolytic cleavage reaction at the membrane interface, designed for the real-time, single-molecule kinetic analysis. The interactions between the enzyme and substrate near the membrane are examined under different classic scenarios. Our findings reveal that while the membrane environment significantly enhances enzymatic activity, it also imposes diffusion limitations that reduce this activity over time. By adjusting the enzyme’s membrane affinity to an intermediate level, we enable the enzyme to "hop" on the membrane surface, overcoming these diffusion constraints and sustaining high enzymatic activity with faster kinetics. These results provide critical insights into the role of the cell membrane in regulating biochemical reactions and can be broadly applied to other membrane-associated interactions.

## Introduction

Plasma membrane is the key component in the cell. It serves as a physical barrier that separates the cell from the extracellular space while maintaining two distinct reaction systems of different dimensions. In addition to the role of three-dimensional (3D) compartments in the cell, cell membrane serves as a two-dimensional (2D) platform for the diverse cellular activities involved in the communication across the membrane. In addition to the membrane proteins, many biomolecules around the cell membrane directly or indirectly participate the critical processes such as the signal transduction, transportation and cell-cell recognition (1–7). However, in terms of the signal transduction, not only does the signaling pathway involving membrane proteins take place on the membrane, but also the subsequent steps occur on the membrane via the recruitment of the cytoplasmic proteins to the surface of the membrane. Why does the cell membrane recruit multiple downstream cytoplasmic signaling proteins specifically to the membrane instead of leaving most of the signaling relay in the cytosol? For example, the classical receptor tyrosine kinase, epidermal growth factor receptor (EGFR), localized on the cell membrane regulates the survival and proliferation of the cell through the mitogen-activated protein kinase (MAPK) pathway. Upon binding of ligands, i.e. epidermal growth factor (EGF) (8), to the extracellular domain of EGFR, EGFR undergoes dimerization, leading to phosphorylation of tyrosine residues in its cytoplasmic tail (9–13). The activated EGFR recruits Grb2 to the cell membrane through the association between phosphotyrosine (pTyr) and the Grb2 SH2 domain (14, 15). Right after the recruitment of Grb2, the guanine exchange factor, SOS, is subsequently recruited from the cytosol into the signaling chain through a series of activation steps. The activation of the membrane-associated Ras protein occurs at the molecular assembly which links EGFR-associated Grb2 with Ras via SOS at the membrane surface (16). Another type of membrane receptor, G-protein coupled receptor (GPCR), also forms the signaling complex at the membrane surface. Upon agonist binding, GPCR undergoes conformational rearrangement, that increases the association between receptor and G-protein. Activated receptor subsequently recruits the heterotrimeric guanine nucleotide binding proteins (G-protein) composed of one Ga, one Gb, and one Gc subunit (17–22), forming the activated agonist-bound GPCR-G protein complex (23). From the examples above, it is obviously that the membrane surface serves as a hub for ligands, receptors, and downstream proteins to form a signaling complex (24, 25). The 2D structure of the membrane can provide additional advantages such as the higher local concentration after the recruitment to the membrane (2, 16, 26), the low degree of freedom for the higher reactivity due to the constrain on the membrane (27), the protein condensate working as a reaction center facilitating downstream signals (28–32) and the ability to shape the conformational energy landscape of membrane proteins to drive reactions (33). Taken together, a 2D platform like the cell membrane is essential for the biological activities in the cell.

The advantages of 2D system for the biochemical reactions in the cell has been discussed for decades. For example, Adam and Delbrück proposed that the reduction in dimensionality can accelerate the rate of biological reactions by enhancing the association between two components related to membrane binding and membrane-bound species (34). In recent years, there have also been theoretical and experimental studies indicating that membrane localization can accelerate association (35, 36). When two proteins are bound to the membrane, not only is lateral movement restricted, but the rotational degree of freedom of individual molecules is also reduced. As a result, when two membrane-bound proteins interact, the entropic cost is significantly lower compared to proteins with high degrees of freedom in the solution (2, 27). Pólya’s theorem shows that when a molecule is bound to a 2D surface, there is a higher chance of encountering another molecule on the surface through a random walk compared to the collision in 3D solution (37). In fact, the membrane localization of the cytosolic enzyme through binding is not entirely a positive factor for the enhancement of the reaction rates. The cell membrane would reduce the diffusion of the enzyme binding to it by nearly 100-fold, lowering the corresponding encounter rates of the proteins on the membrane by about 3 to 30-fold (2). The diffusion-limited kinetics would dictate the reaction of the proteins anchored on the membrane. On the other hand, the membrane confining molecules to the 2D surface can increase the local concentration of proteins by a factor of more than 600 (2, 27, 38). Due to the high local concentration of the molecules on the membrane, the membrane is able to facilitate the association between two interacting proteins, a process often impossible in a solution (39). In addition to the reduction of the degree of freedom due to the confinement on the membrane enhancing the reaction rates (2, 27), the heterogeneous catalytic effect of the cell membrane can be further enhanced via the surface-confinement effect.

Cell membrane as the essential feature in the cell provides the heterogeneous catalytic environment to speed up the biological reactions. In addition to the catalytic effect of the *in vivo* membrane systems, the concept of the surface confinement has been applied to the *in vitro* systems (40, 41). Studies show that the biological reactions are facilitated by the attachment of the soluble enzymes to the solid support providing the enhanced stability of the enzymes (42, 43). The enzymes with the enhanced stability through the surface confinement can also reach higher regio- and stereoselectivity (44–46). Sophisticated systems involving enzymatic cascade reactions reveal the critical role of the spatial control on the enzymes. The enzymatic activity is mediated by the averaged distance among the enzymes (41, 47–49). Besides the surface confinement and the direct spatial control of the enzyme, more and more related mediating effects on the reaction at the membrane surface such as the crowding effect are addressed. The crowding effect is an important dynamic spatial control in most biological reactions (50–54). In biology, high concentration of macromolecules occupies a large amount of space in the fluid media. The combined concentration of various macromolecules can reach up to 400 mg/ml in cells (55, 56). Therefore, most cellular activities proceed under the crowding condition. For example, previous studies have shown that the crowding effect can facilitate the protein folding (57–59), affect conformational stability (60, 61), and induce phase separation in the aqueous solution (62–65). Moreover, the crowding effect can affect the behavior of the molecule resulting in slower diffusion (66–68), the higher tendency of association with other molecules (69–75) and change the reaction rates and equilibria accordingly. The crowding effect is also expected on the cell membrane as well given that the membrane proteins occupy approximately 30-50% of the surface area on the membrane (76, 77). Furthermore, the lipid composition of the membrane can affect the local environment and the physical properties of the membrane modulating the reaction kinetics on the surface. For example, the accumulation of cholesterol in the membrane slows down the lipid lateral diffusion (78) and lowers the activity of the kinase about 3.5 folds when the bilayer contains 1% cholesterol (79). The shape of the lipid molecule determined by the relative size of the head group and hydrophobic tails of lipids can induce the spontaneous membrane curvature enhancing the interaction among the proteins (80, 81). Cholesterol, sphingolipids, and glycolipids are regarded as the key lipid components of the lipid raft, those specialized microdomains of the membrane are also essential for the cellular signal transduction (82–87). In sum, the characteristic properties of the membrane, such as the low dimensionality, high structural complexity, complicated lipid composition, fluidity across the 2D surface, etc., play the key roles on the regulation of the biochemical processes in the cell.

In this work, we systematically interrogate the biological reaction mediated by the 2D membrane environment using Tobacco Etch Virus (TEV) proteolytic cleavage reaction as the model reaction. We anchored the substrate of the TEV protease on the supported lipid bilayer (SLB) and study the kinetics of the proteolytic cleavage reaction when TEV protease is recruited on the membrane. The enhanced green fluorescence protein (eGFP) is attached to the substrate of the TEV protease as the probe to track the kinetics of the TEV proteolytic cleavage reaction. The eGFP fluorescence from the membrane surface is monitored by the total internal reflection fluorescence (TIRF) microscopy to read out the kinetics of TEV proteolytic reaction on the membrane in real time. The fluorescence intensity is further converted into number of molecules through a series of delicate calibration processes allowing the enzymatic reaction on the membrane to be studied at the single-molecule level. Here we see that the biological reaction could be facilitated by the 2D membrane environment. Our data shows that the activity of TEV protease is vastly enhanced at the beginning compared to the reaction in solution and the reaction with substrate on the membrane and enzyme in the solution. However, the activity of TEV protease decreases dramatically as it depletes the nearby substrates soon after the recruitment to the membrane indicating the diffusion limited scenario of the enzymatic reaction. The membrane affinity of TEV protease is further modulated from high to none through the length of His tag with the presence of imidazole creating the scenarios from the permanently membrane-bounded, surface-hopping and collision-type enzymatic reactions. The TEV protease exhibits the highest activity if the reaction is proceeded via the surface hopping of TEV protease. Our study identifies the role of the membrane for the reaction at the membrane interface. To efficiently utilize the membrane environment to speed up the reaction on the membrane, the enzyme should have the moderate affinity to the membrane to break the diffusion-limited condition. The enzyme can locally deplete the substates several times via multiple recruitments to the membrane. In the end, our work implies the necessity of the adapter protein such as Grb2 using SH2 and SH3 domains to bridge the phosphotyrosine of RTK to the proline-rich domain of GEF protein, i.e. SOS. The compromised affinity due to the synergy of two different associations via SH2 and SH3 results in the effective activation of G protein in a short period of time.

## Result

To study the biological reaction on the membrane under different scenarios, we reconstructed the TEV proteolytic cleavage reaction on SLB as depicted in Figure 1B. The substrate of TEV protease has a TEV cleavage sequence after a His6 tag at the N-terminus and a GFP at the C-terminus. GFP serves as the probe to monitor the cleavage process. Once the TEV cleavage site is cut, the GFP is released into the bulk solution in the background. Given that the fluorescence of GFP in the bulk solution is not collected by the total internal reflection fluorescence (TIRF) microscopy, the cleaved substrate can be determined via the decrease of the GFP fluorescence on the membrane to extract the kinetics of the TEV proteolytic cleavage reaction. The conversion between the GFP fluorescence on the membrane and the corresponding number of GFP molecules is done by a series of calibrations (see Figure S1 in the supporting information). In short, the EMCCD with the single-photon sensitivity is used to collect the single-molecule images. The number of the molecules on the SLB and the corresponding total fluorescence intensities of the single-molecule images are linearly correlated (See Figure S1). The ratios of the total fluorescence intensities across two different settings of EMCCD from the images with the density of the substrate at or above the single-molecule are used to convert the total fluorescence intensities into the exact number of substrates per square micrometer on the SLB. The substrates of TEV protease are anchored on the SLB with 8 mole% nickel nitrilotriacetic acid (Ni-NTA) lipid through histag chemistry. The fluorescence recovery after photobleaching (FRAP) experiments confirm the 2D Brownian motion of TEV substrate on the lipid bilayer (see Figure S2). TEV was labeled by Alexa Fluor 647 using NHS ester chemistry. The histag of TEV allows TEV to be recruited on the SLB after the addition to the bulk solution above the SLB, and the fluorescence of TEV measured by TIRF is directly converted into the number of TEV molecules per square micron like the substrate tagged with GFP.

**Figure 1.**
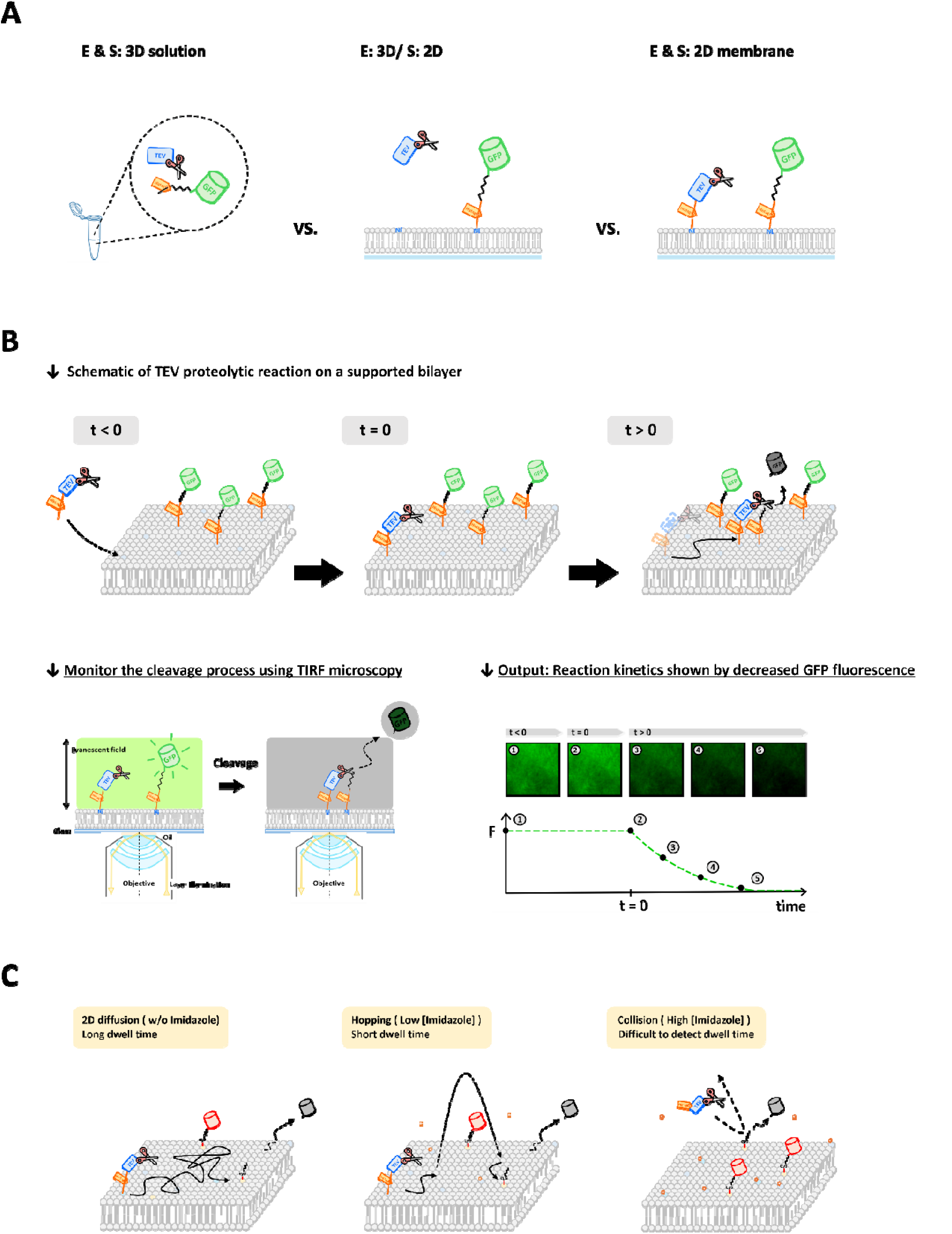
(A) Comparison of TEV proteolytic reaction in different dimensions. (B) Schematic of TEV proteolytic reaction reconstruction on the supported bilayer. His-tagged GFP is anchored on a supported bilayer containing 8% Ni-NTA DOGS. His-tagged GFP has a TEV cleavage site that connects the His-tag and GFP. His-tagged TEV is recruited to the supported membrane by His-tag chemistry and undergoes lateral diffusion. TEV recognizes and cleaves the cutting site, releasing GFP from the SLBs. Enzyme activity for the cleavage of the peptide bond is determined by a decrease in GFP fluorescence on the surface. (C) Modulation of His4-TEV motion type by adjusting the affinity between the His-tag and Ni-NTA lipids.

The proteolytic cleavage reaction begins right after the recruitment of TEV on the SLB. In Figure 2A, the number of TEV on the SLB increases rapidly at the beginning and reaches the plateau after 5 minutes mimicking the signal transduction right after the activation of the receptor. Given the TEV is not autoinhibited, the beginning of the recruitment is treated as the time zero of the proteolytic cleavage kinetics. TEV laterally mobile on the 2D surface encounters the surrounding GFP tagged substrates and processes the cleavage of the TEV cutting site on GFP. The release of GFP from the SLB is reflected by the decrease in GFP fluorescence on the surface during TEV cleavage. The control experiment without TEV shows the negligible fluorescence decay (see Figure S3A). The fluorescence intensity on the SLB is monitored every 5 seconds by capturing the TIRF fluorescence images of GFP and TEV. The fluorescence intensities at each time point are further converted into the number of the GFP substrates. The decrease of the GFP substrates between two adjacent time points shows the number of the cleaved GFP substrates by TEV. The ratio of the cleaved GFP substrates to the instantaneous quantity of TEV on the SLB is further calculated to obtain the real-time TEV activity.

**Figure 2.**
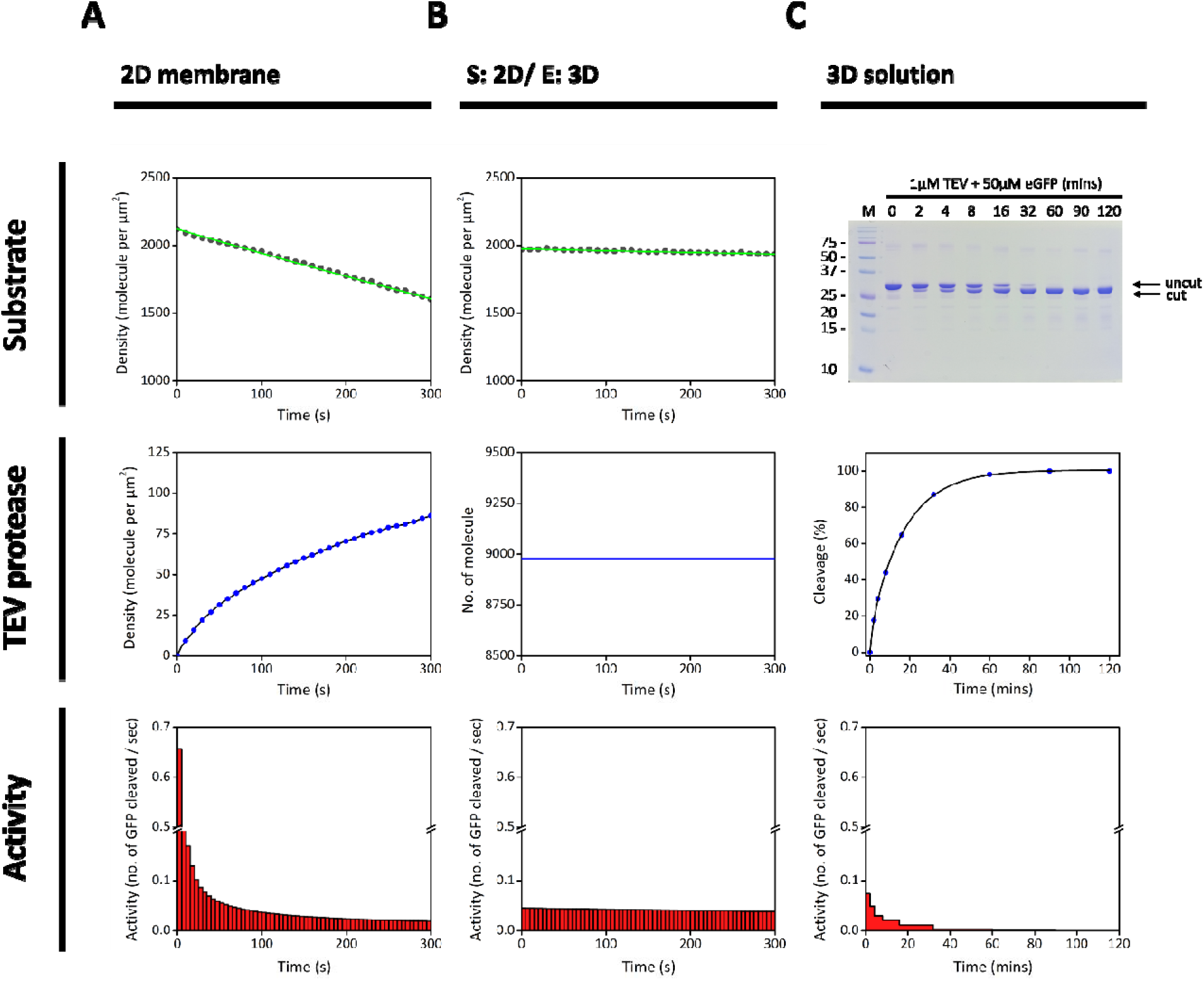
2D surface enhances the reaction rate at first by higher local concentration. (A) TEV proteolytic reaction on membrane. Fluorescently labeled His7-TEV is recruited to supported membrane (Blue line). GFP anchored on the membrane is released from the surface due to proteolytic cleavage (Green line). (B) TEV proteolytic reaction between 2D and 3D system. Cleavage may occur when TEV in solution collides with substrates anchored on the membrane. The number of non-His tagged TEV distributed within a space near the membrane with a thickness of one hundred nanometers is 9000 within the observed area. (C) TEV proteolytic reaction in solution. TEV (final 1µM) is mixed with GFP (final 50µM) in 0.1M PB buffer solution. SDS-PAGE is used to track the kinetics of the TEV proteolytic cleavage reaction. Enzyme activity is determined by calculating the area ratio of cleaved substrate to uncleaved substrate.

In Figure 3A and 3B, the surface densities of GFP substrate on the bilayer are prepared at around 2000 and 1000 molecules/µm^2^ respectively. The recruitments of TEV reaching different surface densities at 100, 50, and 20 molecules/µm^2^ are prepared by incubated the GFP substrate anchored on the SLB with the different concentrations of TEV solutions. The sample with the higher amount of TEV recruited on the membrane leads to the faster decrease in the density of GFP substrates. Additionally, when the ratio of the GFP substrate to TEV molecules/µm^2^ is large, the initial activity of TEV has the higher number with the TEV activity of 0.66 molecules per second for the ratio of 20, the TEV activity of 3.08 molecules per second for the ratio of 40, and with TEV activity of 4.66 molecules per second for the ratio of 100 indicating that GFP substrate becomes in excess. TEV competes for the limited surrounding GFP substrates. The real-time TEV activity of each condition decreases quickly implying that the depletion of GFP substrates occurs soon after the recruitment of TEV on the SLB and the proteolytic cleavage reaction is limited by the diffusion on the SLB.

**Figure 3.**
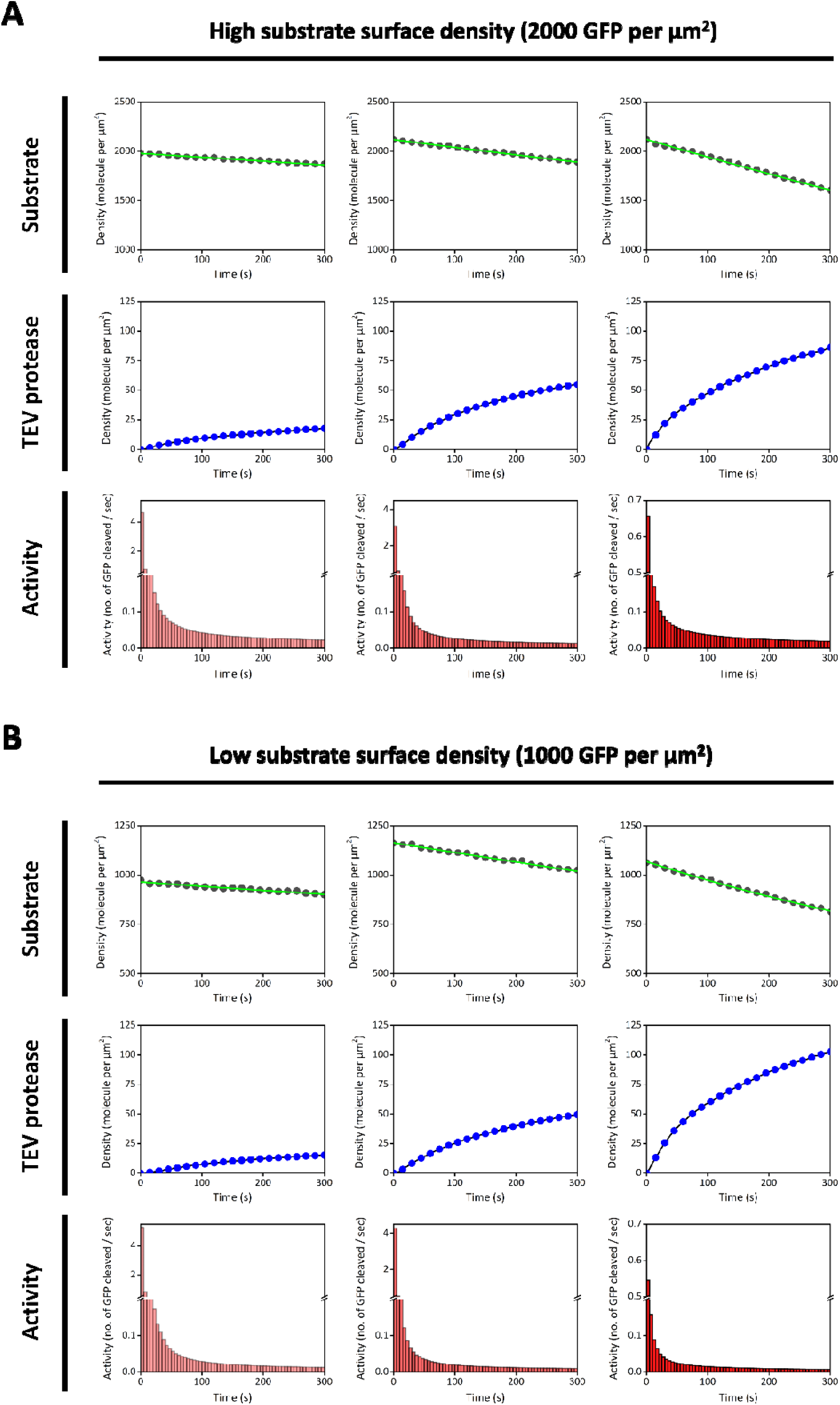
The TEV activity is enzyme surface density dependent. During the reaction, the number of GFP decreases due to proteolytic cleavage by TEV protease (Green line). After injection, His7-TEV is recruited to the membrane through His-tag chemistry (Blue line). Time course of TEV activity is shown (Red bar). Each bar represents the average number of substrates that one TEV molecule can cleave per second. The activity of TEV depends on the surface density of enzymes and shows competition for substrates among enzymes. The data sets demonstrate that more TEV proteases are recruited to the membrane, lower enzyme activity measured due to fewer substrates being distributed to each enzyme. The surface density is controlled at about 2000 molecules per µm^2^. A similar density trend in activity was observed in lower substrate surface density (1000 molecules per µm^2^).

To explore the highest turnover number of TEV on the SLB without the competition for the limited number of GFP substrates, we constructed the proteolytic reaction with TEV at the single molecule level (see Figure 4). The number of recruited TEV on the SLB is controlled between 400 and 500 in the observed area (55 micron by 55 micron) through the incubation of 0.05 nM TEV solution on the SLB anchored by GFP substrate. Compared to the bulk experiments in Figure 3, the dependence of the TEV activity on the ratio between GFP and TEV holds with TEV at the single molecule level. The ratio between GFP and TEV is much higher than the ratio in the bulk experiments resulting in the initial TEV activity larger than ∼80 molecules per second. Moreover, the fast drops of TEV activities like the TEV activity of the bulk experiments in Figure 3 are also observed implying that the similar diffusion limit is reached in about few seconds.

**Figure 4.**
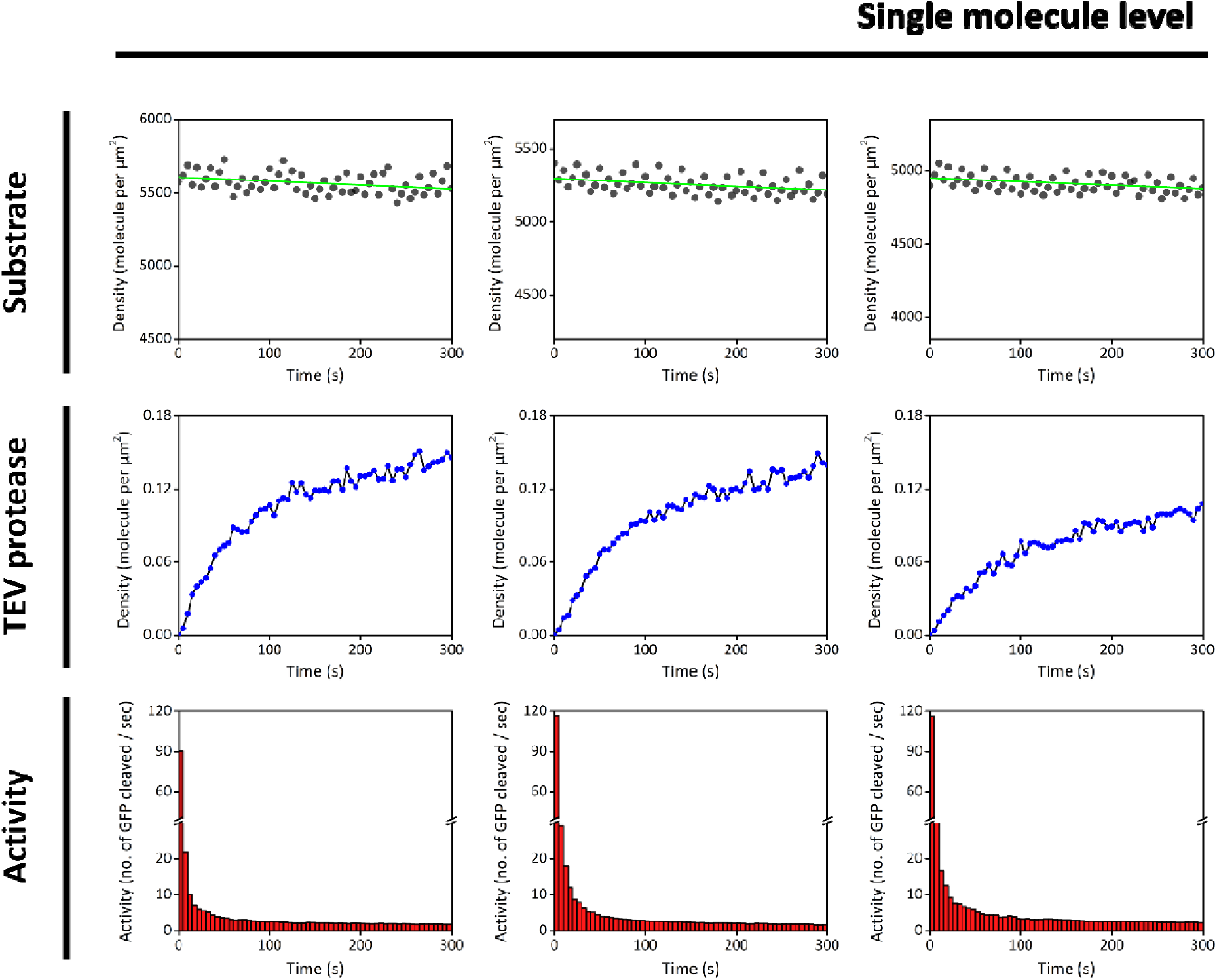
TEV activity is increased by 2D membrane without competition between enzymes. The activity of TEV is dramatically enhanced when both TEV and its substrates are anchored to the membrane, without competition between enzymes for substrates. In this experiment, there is a large excess of substrates compared to TEV to prevent competition between enzymes for substrates. The surface density of GFP tagged substrates on the membrane is controlled at 5000 molecules per µm^2^ (Green line), while the number of TEV recruited to the membrane is controlled at 400-500 (surface density of approximately 0.12 molecules per µm^2^) within the observed area (Blue line). The initial activity of TEV is boosted to a couple of hundred (Red bar).

Given that TEV and GFP substrate are both anchored on the SLB via Histag chemistry, the potential competition for the Ni-NTA binding sites can lead to the dissociation of GFP substrate from the SLB and result in positive bias on TEV activity. To verify that the fluorescence decrease of GFP is only due to TEV cleavage rather than competition for Ni-NTA binding sites, we mutated the cystine to alanine at residue 151 located at the catalytic triad of TEV. The catalytic capability of C151A TEV mutant (TEV^C151A^) has been reported to be impaired, but the structure of TEV is not disrupted by this mutation (88). The activity of TEV^C151A^ is (further examined and shown by SDS-PAGE in Figure S5. TEV^C151A^ (28 kDa) in the SDS-PAGE is located at the position of higher molecular weight (71 kDa) comparing to the standards indicating that the Maltose binding protein (MBP) fusion tag (43 kDa) is retained after the protein expression. Given that MBP tag is fused with TEV for the higher protein yield and self-cleaved by TEV at the TEV cutting site between MBP and TEV, the result above confirms that TEV^C151A^ is indeed non-functional. However, the remaining MBP fusion tag blocks exposure of the His7 tag located between TEV and MBP to the Ni-NTA lipid and inhibits the recruitment of TEV to the SLB. Therefore, we removed MBP from DNA sequence and used His7 tag– TEV^C151A^ as the control in our experiment. Under the same condition, TEV^C151A^ is introduced to the SLB anchored by GFP substrates, and the fluorescence of GFP remained nearly the same during the recruitment of TEV^C151A^ confirming that the decrease in GFP fluorescence during the recruitment of the wildtype TEV is due to the proteolytic cleavage by TEV (see Figure S4).

In the solution-type experiments (3D), TEV and GFP substrate of the proteolytic cleavage reaction are prepared at the micromolar level (1 µM and 50 µM) with a molar ratio of TEV to GFP at 1:50 in PB buffer (pH 8) which are about two orders of the magnitude higher than the concentrations of TEV and GFP substrate solution used in 2D experiments. The kinetics of cleavage reaction is monitored at different time points by collecting and flash freezing 2µl of the reaction mixture for SDS-PAGE analysis. The SDS-PAGE analysis in Figure 2C shows that most of the GFP substrates are cleaved by TEV shown by the band corresponding to the lower molecular weight in 90 minutes. The percentage of the cleaved substrates quantified by ImageJ is used to plot the reaction kinetics. The corresponding real-time TEV activity in the aqueous solution suggests that in average a small number (0.074) of GFP substrates are cleaved by one TEV in the first 120 seconds. In 2D experiment, the average TEV activity in first 120s is about 0.093 molecules per second (under condition with reactant concentration lowers by 1-2 orders of magnitude than in 3D solution). To further interrogate the role of the membrane on the proteolytic cleavage reaction at the interface, we separate the enzyme and the substrate into different dimensionality. The membrane affinity of TEV is shut off by the removal of its Histag and prepared in the bulk solution above the SLB. To do so, Factor Xa cutting site is inserted between TEV and the His7 tag. The non-Histagged TEV is generated after the removal of the N-terminal His7 tag via Factor Xa cleavage during the purification. The GFP substrate is anchored on the SLB like the 2D experiment through Histag chemistry. The proteolytic cleavage reaction is now mainly through collision between the enzyme and the substrate. The corresponding reaction rate is dictated by the frequency of collisions of TEV from the bulk solution. The number of TEV distributed at a height of 100nm from the surface of the solution is estimated based on the injection concentration. During the reaction process, the density of the GFP substrate only slightly decreases. The TEV activity is nearly constant, approximately at 0.045 molecules per second (see Figure 2B).

In a 2D experiment, we observed a rapid decrease in TEV activity 5 seconds after the reaction started due to reduced diffusion. In biology, signaling cascades are achieved by low-affinity (1-100 nM) and readily reversible interactions between cytoplasmic adaptor proteins and membrane receptors. After recruitment, the adaptor protein is not confined to the plasma membrane for long but sequentially binds to multiple receptors through a protein hopping mechanism. In order to further explore, we introduced imidazole to modulate the His-tag/Ni-NTA affinity, further controlling the motion type of molecules on the membrane. Four different SLB environments are prepared with imidazole concentrations of 0 mM, 10 mM, 20 mM, and 40 mM. Injection of 0.05 nM His4-TEV sample corresponds to TBS buffer with imidazole concentrations. With a total internal reflection fluorescence (TIRF) microscopy, we can monitor the dwell time of individual His4-TEV molecules by tracking the molecules from their recruitment to the SLB via His-tag chemistry to their disappearance due to competition with imidazole and leaving the TIR illumination field. Histograms of the dwell time distribution for His4-TEV are shown in Figure 5. As expected, the dwell time of His4-TEV is shorter when there is a higher concentration of imidazole in the system.

**Figure 5.**
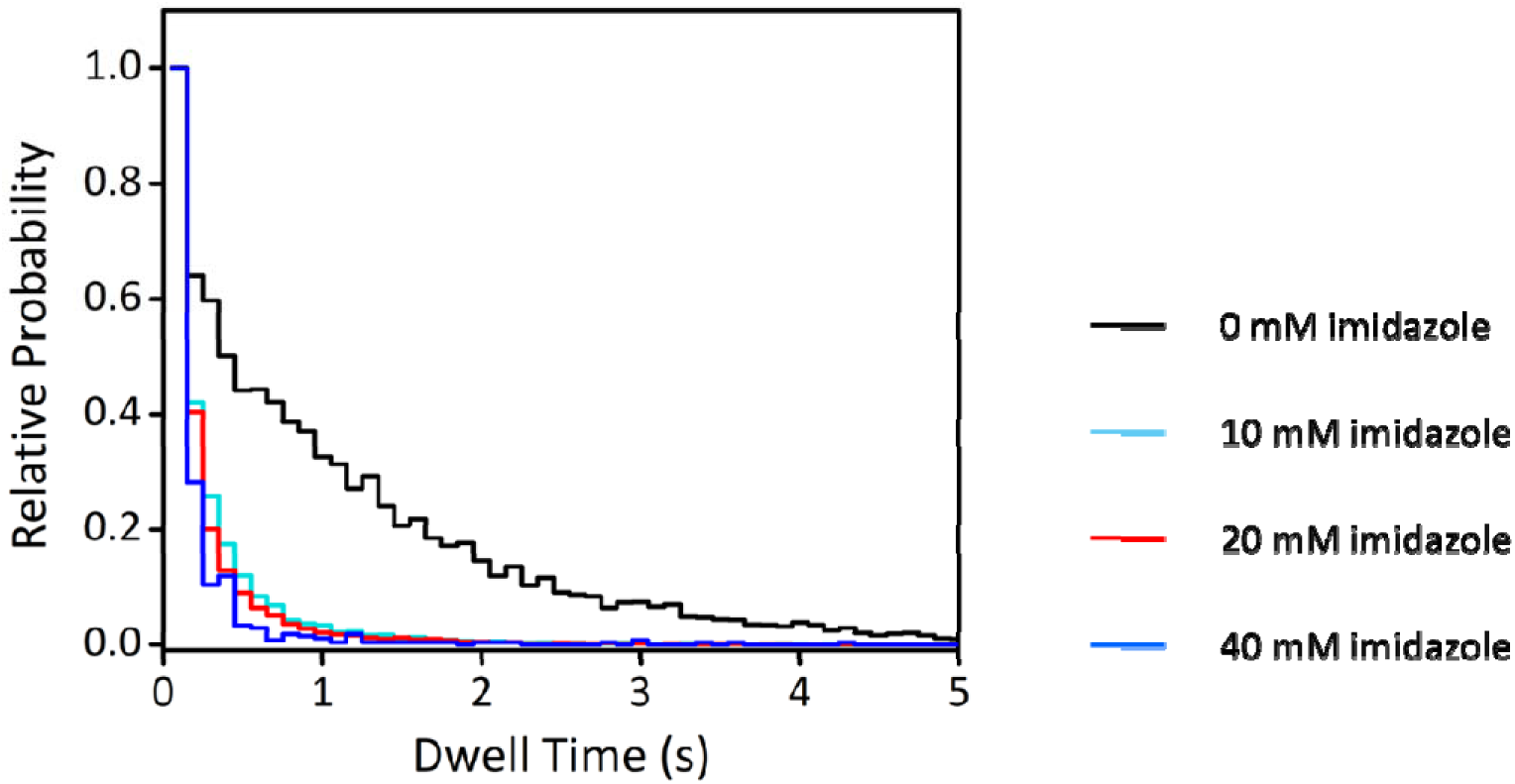
Modulation of His4-TEV dwell time via imidazole concentration adjustment. Dwell time distribution for His4-TEV under four conditions with imidazole concentrations of 0 mM (black), 10 mM (cyan), 20 mM (red), and 40 mM (blue). Imidazole competes with the His-tag for binding to the Ni-NTA lipid, causing His4-TEV to leave the membrane and shortening its dwell time. The histogram shows tails sparsely populated with states extending toward longer dwell times when there is no imidazole in the system; as the imidazole concentration increases, the distribution becomes more concentrated around shorter dwell times.

After confirming that imidazole can modulate the motion type of molecules on the membrane, we introduced imidazole to the reconstitution of proteolytic cleavage on SLB to further explore how the motion mode of molecules on the membrane affects reaction rates. The introduction of imidazole would cause the GFP substrate to become unstable since it is also anchored to the SLB through His-tag chemistry. Hence, we inserted a cysteine residue at the N-terminus of the substrate and replaced GFP with mCherry as the probe to monitor the cleavage process. The mCherry substrate is anchored to the membrane by coupling the cysteine at the N-terminus to maleimide lipids (PE MCC), as depicted schematically in Figure 1C. Because mCherry itself does not have cysteine residue in the protein sequence, the cysteine is the only anchoring point between the substrate and the bilayer. We mimicked three motion types of His4- TEV: 2D diffusion, hop diffusion, and collision, corresponding to imidazole concentrations of 0 mM, 40 mM, and 200 mM, respectively (see Figure 1C). In the absence of imidazole ([IM] = 0 mM), the interaction between the His-tag and Ni-NTA lipid is strong. His4-TEV is recruited to the membrane via His-tag chemistry and undergoes 2D diffusion. This 2D surface initially enhances TEV activity to around 0.78, followed by a rapid decay due to the diffusion-limited condition on the membrane. The introduction of a low concentration of imidazole ([IM] = 40 mM) reduces the affinity between His4-TEV and the membrane surface containing 4 mole% Ni-NTA lipids. This reduction in affinity causes TEV to undergo hopping movement on the SLB. The motions of TEV include the recruitment to the SLB, 2D diffusion on the SLB, dissociation from the SLB, and rebinding to the membrane, which expands the effective range of TEV during the reaction and overcomes the slow diffusion exerted by the membrane. The kinetic traces are faster and the corresponding enzymatic activities are much higher compared to the case without imidazole. At the high concentration of imidazole ([IM] = 200 mM), His4-TEV cannot be recruited to the membrane. TEV interacts with the substrate only through collisions, leading to relatively low activity (see Figure 6).

**Figure 6.**
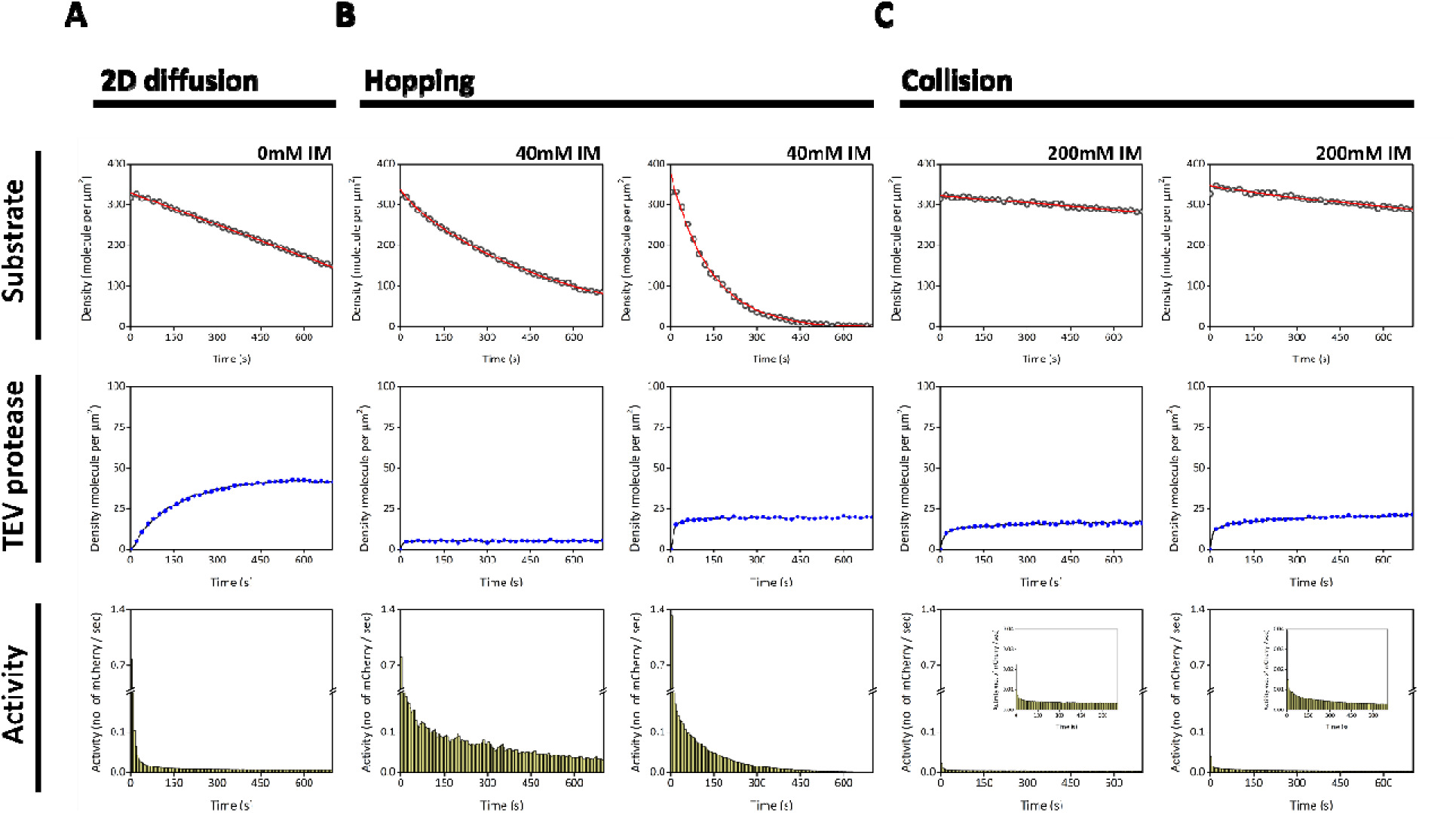
Protein hopping can overcome localized movement caused by the reduction of diffusion when a molecule is tethered to the membrane. Cys-TEVcut-mCherry is anchored on the membrane by maleimide chemistry (with a surface density controlled at about 350 molecules per µm^2^). The motion type of His4-TEV is modulated by the affinity between the His-tag and Ni-NTA lipid. (A) Without imidazole ([IM]=0mM) in the system, the interaction between the His-tag and Ni-NTA lipid is strong. His4-TEV is recruited to the membrane by His-tag chemistry and undergoes 2D diffusion on the membrane. The 2D surface enhances the initial TEV activity (to about 0.7), followed by a rapid drop and entry into a diffusion-limited state. (B) The introduction of low concentration of imidazole ([IM] = 40mM) reduces the affinity between His4-TEV and the membrane surface containing 4% Ni-NTA lipid. This reduction in affinity causes TEV to undergo hopping movement on the supported membrane. The movement processes of TEV involve recruitment, 2D diffusion, hopping, and rebinding, which expand the movement range of TEV and are not limited to slow diffusion. TEV can maintain high activity, unaffected by diffusion limitations (on the left). However, in the presence of a large amount of TEV, activity quickly declines due to being reaction-limited (substrates have been completely reacted). (C) His4-TEV is unable to be recruited to the membrane due to the high concentration of imidazole ([IM] = 200mM). TEV reacts with the substrate only through collisions, resulting in relatively low activity.

## Discussion

In this work, the role of the membrane for the biological reaction at the interface is systematically interrogated. The biological kinetics mediated by the 2D membrane has been discussed for decades. Several studies have pioneered the theoretical framework for the biochemistry confined at the interface. Due to the limited number of experimental studies on this topic, our work aims for a clear picture of how the biological reaction mediated by the 2D membrane environment. Through the evolution, the membrane of the cell not only develops as the physical barrier for the polar or charged molecules, but also serves as the reaction center for the membrane proteins. For example, in catalysis it is known that the effective concentration on the heterogeneous surface can be largely enhanced (89, 90). Given that molecules on the membrane are mobile following the 2D Brownian motion, the 2D membrane works as the ideal platform for the reaction among the membrane proteins or the molecules around it. However, studies also show that the diffusion of the molecule affiliated to the membrane is one to two orders of magnitude lower than the diffusion of the molecule in the 3D aqueous solution retarding the reaction kinetics. (references such as 16 27 35 98 99 100) More and more factors mediating the reaction kinetics on the membrane are identified such as rotational degrees of freedom and thermodynamic stability of protein complex (27). It is widely hypothesized that the enhanced local concentration by the membrane can speed up the reaction rate due to the higher encounter rate. (2, 16) Recently, Huang *et al.* has shown that the association is 22- to 33-fold faster at the membrane than in solution for a 10 µm spherical cell by DNA strand-displacement reaction (35). Although the enhancement of the reaction rate achieved by the high local concentration due to the reduction of the system dimensionality is greater than the decrease of the reaction rate caused by the low mobility on the membrane (16, 27, 91). The slow diffusion leads to the localization on the membrane, making the membrane-associated reaction more prone to reach the diffusion limited condition where the number of the substrates around the enzyme is depleted and the replenish of the substrate is limited by the slow diffusion on the membrane. The kinetics on the membrane is clearly dictated by the combination of pros and cons intrinsically coming from the 2D environment of the membrane. Besides the encounter rate, other factors needed to be interrogated experimentally and the real biochemical reaction must be studied directly on the membrane. In this study, we observed that the molecules hopping on the membrane can overcome this limitation. Many biomolecules involved in signal transduction have the medium level of binding affinity such as the micromolar dissociation constant between the SH2 (Src Homology 2) domain of Grb2 and the phosphotyrosine and between GPCR and G-protein (92, 93). Molecules hopping on the membrane have moderate affinity to the membrane. Studies have shown that the adapter proteins can reversibly bind to the receptors anchored on the membrane (94, 95). For example, Grb2 is recruited to the membrane via the binding to phoshpotyrosines on EGFR c-tail. Given the micromolar dissociation constant between Grb2 and EGFR, Grb2 can hop among the phosphotyrosines across multiple receptors increasing the encounter of the downstream protein and resulting in the signal propagation. (96, 97) A model reaction of TEV proteolytic cleavage reaction without additional modulation on the enzymatic activity via allostery or coenzyme is *in-vitro* reconstituted on the SLB to study the biochemical kinetics directly on the membrane. In the first part, both the enzyme (TEV) and the substrate (eGFP tagged TEV cutting sequence) are prepared on the SLB or in the bulk solution resulting in three scenarios (see Figure 1A). For the enzymatic reaction on the SLB, the TEV proteolytic cleavage of the substrate would result in the dissociation of eGFP fusion tag into the bulk solution above the SLB. The decrease of the fluorescence intensity on the SLB monitored by TIRF microscopy is used to read out the kinetics in the real time (see Figure 1B). From these three scenarios, the scenarios that the enzyme and the substrate both on the SLB or in the solution show the clear kinetics of the proteolytic cleavage reaction (see Figure 2A and 2C) while the concentrations of the enzyme and the substrate used to prepare the experiments are differ in an order of the magnitude in each other. Given the strong affinity to the membrane via the histag chemistry, about 100 nanomolar or lower of the enzyme and the substrate is used to prepare the reaction on the SLB. The averaged reactivity of TEV in 2D environment is calculated to be ∼0.093 no. of substrate per second corresponding to the time window of the first two minutes of the reaction in 3D solution. The reactivities of TEV from these two scenarios now look very similar to each other. It shows that the high affinity to the membrane can compensate for the high concentration required for the proteolytic reaction in the solution. On the contrary, the reaction can barely occur if the enzyme and the substrate are separated into the solution and the membrane (see Figure 2B) showing that the recruitment to the membrane can be used to gate the reactants for a reaction on the membrane while the molecules with the nanomolar concentrations in the cytosol can hardly react with each other or participate the reactions on the membrane through collision.

The kinetics of the TEV proteolytic cleavage reaction on the SLB is further studied in detail (see Figure 3 and 4). In Figure 3, the cleavage reactions on the SLB are reconstituted at two different densities of GFP substrate (2000 and 1000 molecules per μm^2^) using similar sets of TEV protease molecules added in the solution. Both parts of the reconstituted cleavage reactions reveal the similar trends that the more enzymes are added the faster kinetics are recorded and the higher ratio of the substrate to the enzyme gives the higher initial enzyme activity. Similar enzyme activities across the low and median levels of TEV between two substrate densities imply that the enzyme is the limiting reagent while the GFP substrates are in relatively excess like the psudo first-order approximation of the bimolecular reaction. The ratio of the substrate to the enzyme is further enlarged by preparing the enzyme at the single-molecule level to examine the upper limit of the enzyme activity in this reconstituted experiment. In Figure 4, the solution of the enzyme at the picomolar (200-folds of dilution comparing to the experiments in Figure 3) is added to the SLB to obtain the number of the enzymes at the single-molecule level in the imaging field. The initial turnover rate of TEV is determined to be >100 per second which is 20- folds higher than the reactivity of the TEV in the experiments with the lower density of TEV. Provided with that the density of GFP substate is similar to the conditions used in Figure 3, the ratio of the initial reactivities between Figure 4 and Figure 3 (the experiments with the low TEV densities) is about 20 which is much smaller based on the first order dependence of the enzyme to the reaction since a 120-fold decrease of the TEV density is applied to the single-molecule experiments in Figure 4. The large deviation of the initial turnover rate can be attributed to the limitation from the dimensionality of the membrane since the substrate can only access the active site of the enzyme from the plane of the membrane.

Besides the initial turnover rates deviate from the first-order dependence on the density of TEV, the TEV activity with respect to time drops sharply in the kinetic trace shown in Figure 3 and 4. Given that the enzyme and the substrate are both on the SLB, the corresponding concentrations of the enzyme and the substrate are calculated to be 28 µM and 700 µM using the thickness of 5 nm above the SLB like this paper (27) to account for the volume at the interface.

Comparing to the concentrations of the TEV and the substrate used in Figure 2C, the kinetics are expected to be faster under these high concentrations of the enzyme and the substrate calculated above. The observed slow kinetics in Figure 3 and 4 is consistent with the fast decay of the TEV activity. The fast decay of the TEV activity can be further explained by the slow diffusion on the membrane. Usually, the diffusion of the membrane protein on the membrane is about two orders of magnitude less than the diffusion of the soluble protein in the solution. The diffusion coefficient of GFP in the eukaryotic cytoplasm is reported to be 30 µm²/s (98). However, when GFP is localized to the membrane, its diffusion decreases significantly to 0.2 µm²/s, representing a 150-fold reduction, as observed by Thomas A. Leonard *et al.* (27). and the other study (35). Similarly, the diffusion of peripheral membrane-anchored proteins, such as Ras, decreases substantially upon membrane association. For instance, Ras exhibits a diffusion coefficient of 20 µm²/s in the cytoplasm (99), which drops to 1 µm²/s when anchored to the plasma membrane (16, 100). Therefore, the enzyme can quickly cleave the nearby GFP substrates once the enzyme is recruited to the membrane, but the replenishment of the fresh GPF substrates is greatly limited by the slow diffusion on the membrane. The diffusion limit of the enzyme and the substrate exhibited by the membrane retarding the encounter between the enzyme and the substrate right after the direct encounter of the enzyme and the substrate during the recruitment of the enzyme on the membrane results in the fast decay of the enzyme reactivity shown in Figure 3 and 4. This phenomenon can be related to the signaling transduction around the membrane where many peripheral membrane proteins have moderate affinity to the membrane upon activation. For example, SOS protein in MAPK signaling pathway can efficiently activate Ras protein anchored on the membrane without permanent attachment to the membrane (101–103). To further test the idea that the short dwell time on the membrane through moderate affinity to the membrane, the concentration of imidazole is used to modulate the membrane affinity of the enzyme via the histag chemistry. The 2D Brownian motion of the fluorescence labeled TEV at the single-molecule level is measured by TIRF microscopy and is used for the analysis of the single-molecule tracking. The histogram of the dwell time on the membrane is plotted in Figure 5 which shows that the dwell time of the TEV molecule decreases as the concentration of imidazole increases. Few single-molecule tracks of TEV molecules on the membrane whose length are shorten by imidazole are shown in SI (Figure S7). The shorter dwell time of TEV due to the modulated histag chemistry can be realized that TEV hops on the membrane surface mimicking the recruitment of the cytosolic enzyme to the membrane with moderate membrane affinity. Since the presence of imidazole would weaken the interaction between the histag and Ni^2+^ NTA lipid, an orthogonal chemistry is required to anchor the TEV substrate on the membrane. In this part, TEV substrate is changed to a similar construct that has the GFP replaced by the mCherry protein. Given the protein sequence of mCherry has no cysteine, one cysteine is added to the c-terminus before the TEV cutting site allowing this TEV substrate to be anchored on the membrane via the maleimide chemistry. The peptide bond of the mCherry at the TEV cutting site can be cleaved by TEV and results in the dissociation of the mCherry part into the bulk solution above the membrane like the GFP substrate. There are three scenarios prepared in this part to understand the role of the hopping mechanism in the biological system (see Figure 1C). In Figure 1C, the first condition is the control part using no imidazole to have both the enzyme and the substrate on the membrane. The second condition has imidazole prepared at the moderate level of the concentration (40 mM) to activate the hopping process of the TEV molecule around the membrane. The third condition have the highest level of imidazole (200 mM) in the bulk solution to prohibit the recruitment of the enzyme to the membrane and limit the interaction between the enzyme and the substrate via the collision of the enzyme from the solution onto the membrane. The density of the molecule in the system is measured and calculated like the previous part, but with a different set of the calibration files exclusively for mCherry protein due to its own characteristic fluorescence. To our surprise, given the similar density of mCherry substrate, the TEV molecule under the hopping condition has faster kinetics than the kinetics from both the enzyme and the substrate anchored on the membrane. With the higher density of the TEV molecule under the hopping condition, almost all mCherry substrates can be cleaved within 10 minutes (see Figure 6). However, the kinetics from the enzyme via the collision under 200 mM imidazole shows the relatively slow kinetics. Notably, the TEV molecule with the hopping mechanism can break the diffusion limit from the membrane and maintain the relatively high enzymatic activity comparing to the TEV permanently anchored on the membrane. Both TEV from the two scenarios have similar initial enzymatic activities, but the enzymatic activity of TEV anchored on the membrane drops quickly due to the slow diffusion on the membrane. The higher density of the hopping TEV on the membrane also indicates the competition of the enzyme for the substrate and results in the fast drops in both the density of mCherry substrate left on the membrane and the corresponding enzymatic activity. The cleavage reaction via the collision of the TEV molecule from the bulk solution under the high concentration of imidazole shows the relatively low kinetics revealing that the high collision frequency of the TEV molecule from the bulk solution is not enough for the reaction on the membrane surface. This observation implies that the affinity to the membrane plays as the switch to trigger the reaction on the membrane which can be one of the mechanisms in the living system regulating the complicated reactions inside the cell.

## Conclusion

In summary, we *in-vitro* reconstituted the proteolytic cleavage reaction on the SLB as the model system to interrogate the role of the membrane mediating the biological reaction at the interface. The dissociation of the fluorescence tag after the proteolytic cleavage reaction allows us to real-time monitor the reaction on the membrane at the single-molecule level. Our work systematically compares the kinetics from different scenarios of the enzyme and the substrate prepared at the interface of the membrane and the bulk solution. From the enzymatic activity of both the enzyme and the substrate anchored on the membrane, it shows that the membrane can increase the interaction between the enzyme and the substrate and give the relatively high enzymatic activity at first. However, the reaction on the membrane is soon subjected to the diffusion limit from the membrane and results in the fast drop of the enzymatic reactivity. We further chemically modulate the membrane affinity of the enzyme to generate the enzyme hopping on the membrane. The reaction kinetics from the hopping enzyme shows the fastest kinetics under the similar condition and can overcome the diffusion limit resulting in the extended decay of the enzymatic activity. Our results provide the key evidence to support the idea that the moderate membrane affinity adopted by nature can fully extract the benefits from the membrane for the high efficiency of the biological reactions at the interface. The hopping mechanism of the enzyme demonstrated in this study can be broadly applied to the interactions among the biomolecules around the membrane regime to better understand how the molecular mechanism behind the scenes is developed through the evolution.

## Contributions

R.B. and C.L. designed research; R.B. performed research; R.B. analyzed data; and R.B. and C.L. wrote the paper.

## Acknowledgements

We would like to thank our undergraduate student, Chun-Chen Lin, for his assistance with DNA cloning and the photobleaching experiments detailed in the Supplementary Information. This work was supported by the grant (111-2113-M-007-030) from the National Science and Technology Council of Taiwan and the grant (113J0064I5) from the Ministry of Education of Taiwan.

## Materials and Methods

### Chemicals

1,2-dioleoyl-sn-glycero-3-phosphocholine (DOPC), 1,2- dioleoyl-sn-glycero-3-[(N-(5-amino-1- carboxypentyl) iminodiacetic acid) succinyl] (Ni^2+^-NTA-DOGS; nickel salt) and 1,2-dioleoyl-sn-glycero-3-phosphoethanolamine-N-[4-(p-maleimidomethyl) cyclohexane-carboxamide] (PE MCC; sodium salt) were purchased from Avanti Polar Lipids. Alexa Fluor 647 NHS ester was purchased from Lumiprobe. Bovine serum albumin (BSA) was purchased from Sigma-Aldrich. Sulfuric acid (H_2_SO_4_) and hydrogen peroxide (H_2_O_2_) were purchased from Honeywell Fluka. Tris-buffered saline (TBS) was purchased from Protech Technology.

### Protein purification

#### GFP tagged substrate

The Green fluorescence protein (GFP) sequence with an N-terminal His6 tag and a TEV protease cleavage site was cloned into p11X plasmid. Transform the plasmid into BL21 (DE3) pLysS *Escherichia coli* using the heat shock method. Transfer the colonies into 1L of Terrific Broth medium and incubate at 37 °C. Once the cell culture concentration reached an optical density at 600 nm (OD600) of approximately 0.6, induced with 1mM isopropyl β-D-1- thiogalactopyranoside (IPTG) and incubated at 37°C for 2 hr. Spin down the cell culture by centrifugation at 6000 rcf for 20 min, then remove the medium. The cell pellet was collected and resuspended in 50 ml Ni-NTA buffer (20mM Tris–HCl, 500mM NaCl, 20mM imidazole, 10% Glycerol, pH 8.0) by vortexing. The resuspended bacteria were lysed by sonication. The cell debris was separated from protein extract by ultracentrifugation at 15000 rcf for 30 min. The supernatant was collected and injected into a HisTrap FF column (GE Healthcare) that had been pre-equilibrium with Ni-NTA buffer. After removing non-target proteins, GFP was eluted by Ni-elution buffer (20mM Tris–HCl, 500mM NaCl, 500mM imidazole, 10% Glycerol, pH 8.0). Concentrate the elution volume to 500µl and exchange to size buffer (0.1 M PB, pH8.0) by an Amicon Ultra Centrifugal Filter Unit (10 kDa molecular weight cutoff [MWCO]; Millipore). Then load the concentrated sample to S75 10/300 column (GE Healthcare) equilibrated with size buffer. The fraction containing GFP was collected, and the purity checked by SDS-PAGE. After determining the concentration, the purified GFP was aliquoted and quickly frozen, then stored in a -80°C freezer.

#### mCherry tagged substrate

pET28 vector containing the sequence of His6-FactorXaCut-Cys- TEVcut-mCherry was transformed into BL21 (DE3) bacteria. Protein expression and purification procedures using a HisTrap FF column (GE Healthcare) and an S75 10/300 column (GE Healthcare) were performed as described in the GFP-tagged substrate purification section. The N-terminal His6 tag was removed by Factor Xa protease, and the cleaved sample was reapplied to the HisTrap FF column. The flow-through was collected, aliquoted, quickly frozen, and stored at -80°C.

#### TEV

pRK508 plasmid containing the TEV^S219V^ mutant fused to an N-terminal His7 and an MBP fusion tag with a tobacco etch virus (TEV) cleaving site was transformed into BL21 (DE3) *E. coli.* The successfully transformed colonies were transferred to 1L of Terrific Broth medium and incubated at 37 °C until the OD600 reached 0.6. Induce the protein expression with 1mM IPTG and grow at 37 °C for 3hr. The culture was centrifuged at 6000 rcf for 20 min, collected pellet and resuspended in 50 ml buffer A (50 mM Hepes, 1 M NaCl, 20 mM imidazole, 10 % glycerol, pH 7.5). The resuspended cells were lysed by sonication, followed by the separation of cell debris from the protein extract through ultracentrifugation at 15000 rcf for 30 min. The supernatant was injected into a HisTrap FF column (GE Healthcare) that had been pre-equilibrium with buffer A. After removing non-target protein, TEV was eluted by buffer C (50 mM Hepes, 50 mM NaCl, 400 mM imidazole, 10 % glycerol, pH 7.5). The elution volume was concentrated to 5 ml and exchanged to buffer B (50 mM Hepes, 50 mM NaCl, 20 mM imidazole,10 % glycerol, pH7.5) by an Amicon Ultra Centrifugal Filter Unit (10 kDa molecular weight cutoff [MWCO]; Millipore). The concentrated sample was loaded onto a HiTrap SP FF column (GE Healthcare) to separate the TEV from non-target proteins by adjusting the concentration of NaCl (altering the ratio between buffer B and buffer D). Collected the elution fraction in which TEV was eluted at approximately 25% buffer D (50 mM Hepes, 400 mM NaCl, 20 mM imidazole, 10% glycerol, pH 7.5). Subsequently, concentrate and exchange to size buffer by an Amicon Ultra Centrifugal Filter Unit (10 kDa MWCO). Then load the concentrated sample to S75 10/300 column (GE Healthcare) equilibrated with size buffer. Collected the fraction containing TEV and measure the protein concentration by UV-Vis. After quick-freezing, store TEV in a -80°C freezer for a later labeling reaction.

### Protein labeling

Protein fluorescence labeling at 1:1 molar ratio. Alexa Fluor 647 NHS ester was dissolved in anhydrous DMSO to prepare a 2 mg/ml dye stock. TEV was concentrated to nine times the volume of the required dye stock to maintain a volume ratio of 9:1 between the protein and the dye. After mixing the protein and the dye, the labeling reaction took place at room temperature for 1 hr followed by 8 hr at 4 °C. The precipitate was separated from labeled sample by ultracentrifugation. Subsequently, the unreacted dye was removed using an Amicon Ultra Centrifugal Filter (10 kDa molecular weight cutoff [MWCO]; Millipore) and the sample was concentrated. The concentrated sample was further purified by size-exclusion column (S75; GE Healthcare). Determine the protein concentration and the labeling efficiency by UV-Vis. Finally, aliquots of the labeled protein were stored at -80 °C after quick-freezing in liquid nitrogen.

### Supported lipid bilayers (SLBs)

Supported lipid bilayers (SLBs) were prepared by vesicle fusion on the glass substrate. Small unilamellar vesicles (SUVs) consisted of the lipid mixture of DOPC and Ni-NTA DOGS at the molar ratio of 92:8 in chloroform. The lipid mixture was evaporated by rotary evaporator for 3 min at 40 °C to remove chloroform and form a dried lipid film followed by N_2_ blow for 15 min to further dry. Add 2 ml of DI H_2_O into the dried lipid film, followed by vortexing and pipetting to resuspend and form vesicles with different sizes. SUVs were formed by dispersing the large vesicle using tip sonication. A glass substrate was treated with a piranha solution that was composed of H_2_SO_4_ and H_2_O_2_ in a volume ratio of 3:1 for 8 min to generate hydroxyl group on the surface (glass coverslips, bottom thickness 170 µm ± 5 µm; Ibidi). Combine the etched glass with the flow chamber (sticky Slide VI 0.4; Ibidi). Mix the SUVs solution and TBS buffer in a 1:1 volume ratio, then inject it into the assembled chamber for 25 min incubation to form SLBs. Incubate 1mg/ml BSA for 15 min to block the defeat in SLBs. GFP was anchored on the Ni-NTA lipid by his-tag-Ni^2+^NTA chemistry for 20 min incubation. The mobility of GFP-associated membrane was examined by fluorescent recovery after photobleaching (FRAP). The density of the protein on the membrane surface was determined by the calibration curve which describes the relation between the average intensity of image and the density of protein within the observed region (see Figure S1 in the supporting information).

### TIRF Microscopy

Imagining was performed utilizing a Nikon Eclipse Ti inverted microscope with a TIRF system and iXon electron-multiplying charged-coupled device camera from Andor Technology. TIRF microscopy with a Nikon 100 × 1.49 NA oil-immersion objective, a TIRF illuminator, a Perfect Focus system, a motorized stage, and a U-N4S four-laser unit as the laser source (Nikon). Equipped with solid-state lasers operating at 488 nm, 561 nm, and 640 nm channels, controlled via a built-in acousto-optic tunable filter. Laser powers were set to 5.2 mW (488 nm), 6.9 mW (561 nm), and 7.8 mW (640 nm), measured with the field aperture fully opened. For filtering, the setup employed the 405/488/561/638 nm Quad TIRF filter set from Chroma Technology Corp. Capture images using the Nikon NIS-Elements software at 5 sec intervals with the exposure time of 20 ms.

### TEV proteolytic reaction

GFP with a TEV cleavage site served as both the reaction substrate and the signal output. Supported membrane contained 8 mole% Ni-NTA lipids to prevent the decrease in GFP fluorescence caused by competition for an anchor lipid between GFP and TEV. Upon the addition of TEV into the supported bilayer with GFP constructs, TEV would be recruited to the membrane and cleave the cutting site, making GFP release from the membrane. The decrease of the fluorescence of GFP from the membrane reflects the process of proteolytic cleavage. The fluorescence of both GFP and Alexa Fluor 647-labeled TEV was tracked at the same time. The average fluorescence intensity of the image was converted to the number of molecules on the surface using a calibration curve to understand the effect of the ratio between substrate and enzyme on the reaction rate. In this study, the injection concentrations of GFP were 250 nM and 125 nM, respectively, to control the number of substrates anchored on the 2D system. One fixed concentration of GFP was used with different levels of recruited TEV (10 nM, 25 nM, and 50 nM). Capture an image at 5 sec intervals and further calculate the real-time TEV activity.

### Imaging Analysis

Mean fluorescence intensity of an image stack over timelapse was analyzed by an ImageJ, Plot Z-axis profile, to obtain the change of fluorescence from GFP on the membrane surface during reaction (or the increase in fluorescence of Alexa Fluor 647-labeled TEV). Single-molecule images of TEV were analyzed by an ImageJ plugin, TrackMate, to count the number of TEV molecules recruited to the membrane. LoG detector was used to localize TEV molecules. The estimated object diameter was set to 0.75 µm. About 400 to 500 molecules were detected per image.

