## Supporting Information for "Enzymatic reactions dictated by the 2D membrane environment"

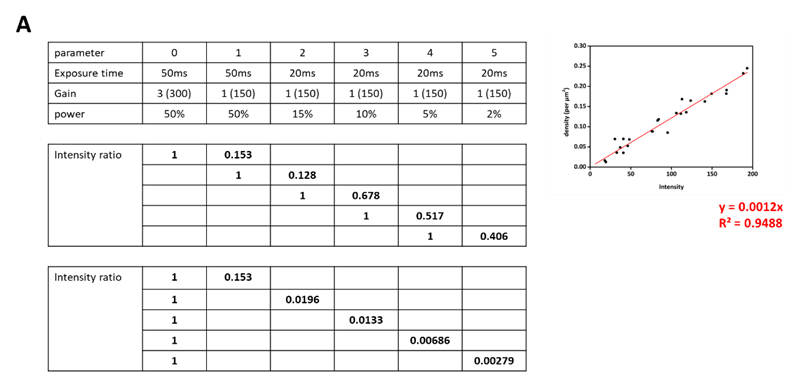

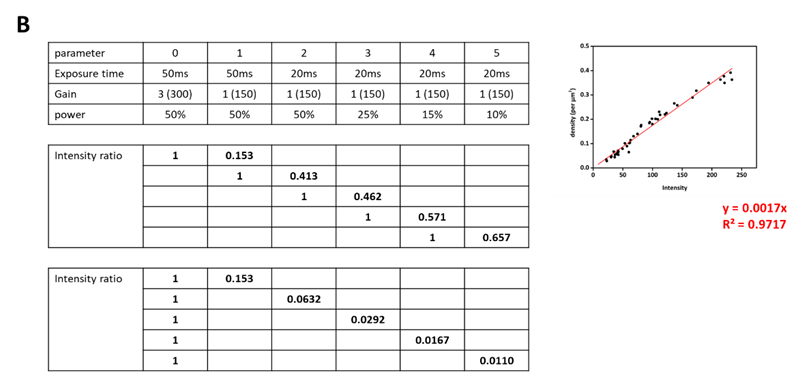

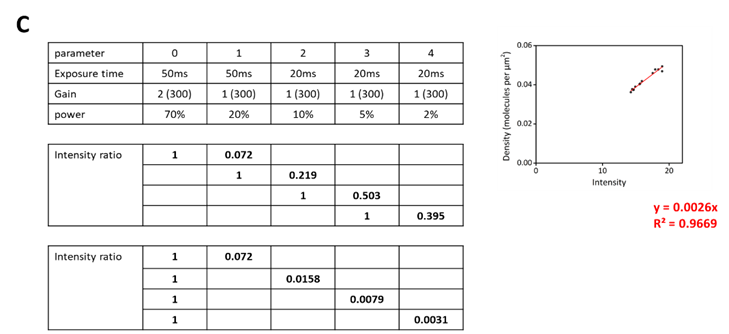

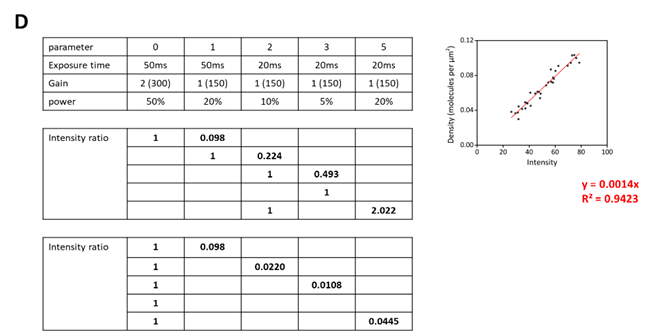

Figure S1. Calibration file for estimating the amount of molecule on the SLB. (A) eGFP. The numbers of the molecules on the SLB and the corresponding total fluorescence intensities of the single-molecule images are linearly correlated. Reconstruction of eGFP on SLB at single-molecule level that can use Image J TrackMate to count the number of molecules. Plot of the relation between the density and fluorescence intensity (right figure). The ratios of the total fluorescence intensities across two different settings of EMCCD from the images with the density of the substrate at or above the single-molecule are used to convert the total fluorescence intensities into the exact number of substrates per squared micrometer on the SLB (left table). (B) His7-TEV. (C) mCherry. (D) His4-TEV.

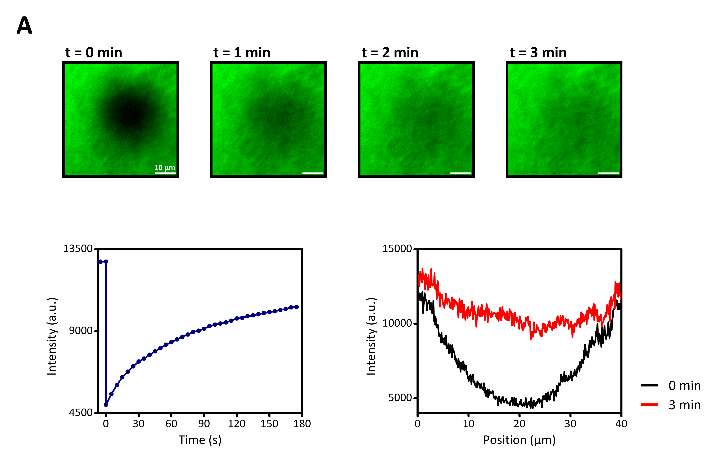

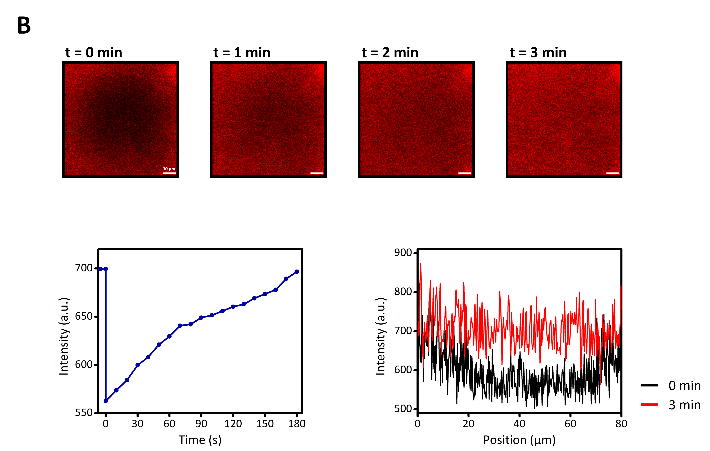

Figure S2. Fluorescence recovery after photobleaching (FRAP) experiment of TEV substrate on supported membranes to show bilayer mobility. (A) FRAP measurements of His6-TEVcut-eGFP on bilayers. After exposure of high intensity light for 1 min, the fluorescent images are taken every 10 sec to monitor the recovery process. Fluorescence recovery > 80% after 3 mins. (B) FRAP measurements of Cys-TEVcut-mCherry on bilayers. After exposure of high intensity light for 1 min, the fluorescent images are taken every 10 sec to monitor the recovery process. Fluorescence recovery > 95% after 3 mins.

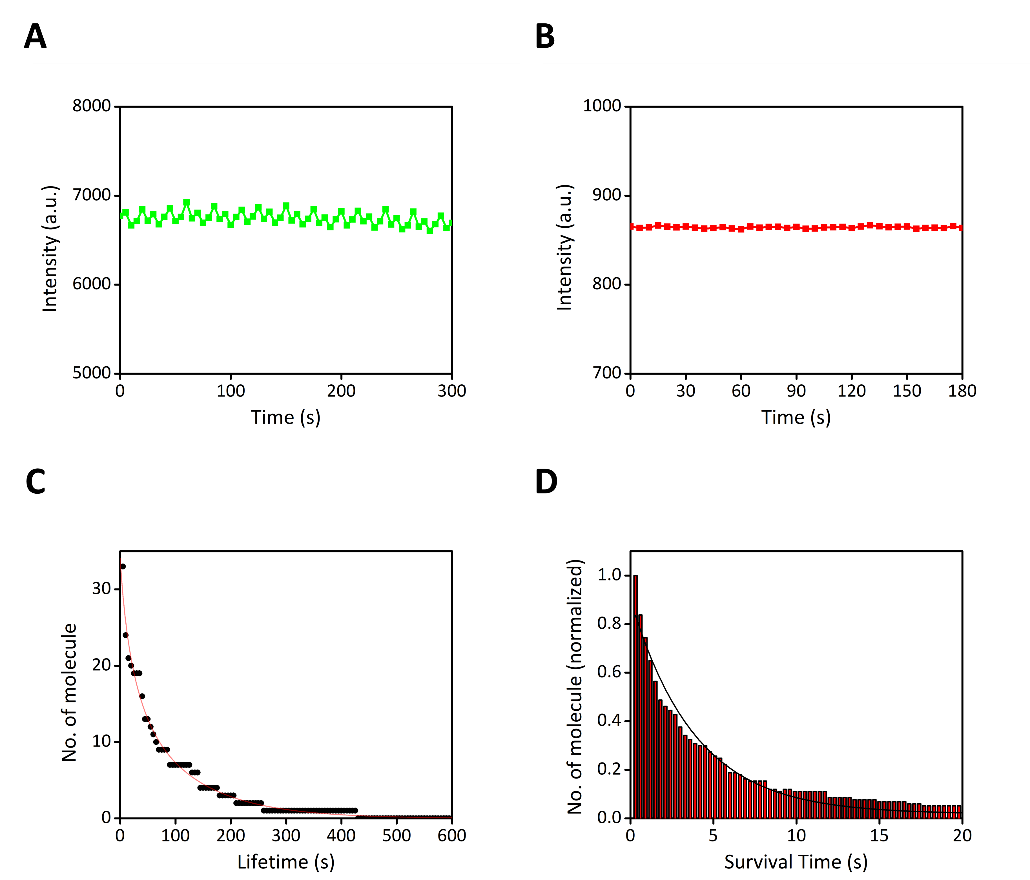

Figure S3. A control experiment to check for photobleaching during imaging. (A) The photobleaching curve of eGFP on bilayers was acquired with an exposure time of 20 ms and laser power set to 2%. Images were captured every 5s. No significant photobleaching of fluorescence was observed during time-lapse imaging. (B) The photobleaching curve of mCherry on bilayers (exposure time: 20 ms, laser power: 2%, imaging per 5sec). No significant photobleaching of fluorescence was observed during time-lapse imaging. (C) The cumulative histogram of the survival times for TEV was fitted with a single exponential function (red line), with a time constant of 7.4 minutes. Alexa Fluor 647-labeled TEV was anchored on an NTA-functionalized PLL-g-PEG surface coupled with Ni^2+^ ions. Images were captured every 5 seconds with an exposure time of 50 ms and laser power set to 30%. In the proteolytic reaction measurement, TEV was mobile. It is inferred that most TEV molecules would not be significantly affected by photobleaching during the measurement period, allowing for accurate quantification. (D) Cumulative histogram of the survival times for TEV. The histogram was fitted with a single exponential function (black line), with a time constant of 3.9 seconds. Alexa Fluor 647-labeled TEV was anchored on an NTA-functionalized PLL-g-PEG surface coupled with Ni^2+^ ions. Images were captured in no delay mode with an exposure time of 50 ms and laser power set to 30%. As a control for photobleaching, to verify the reliability of TEV dwell time measurements, the disappearance of TEV molecules was attributed to leaving the TIR illumination field rather than to photobleaching.

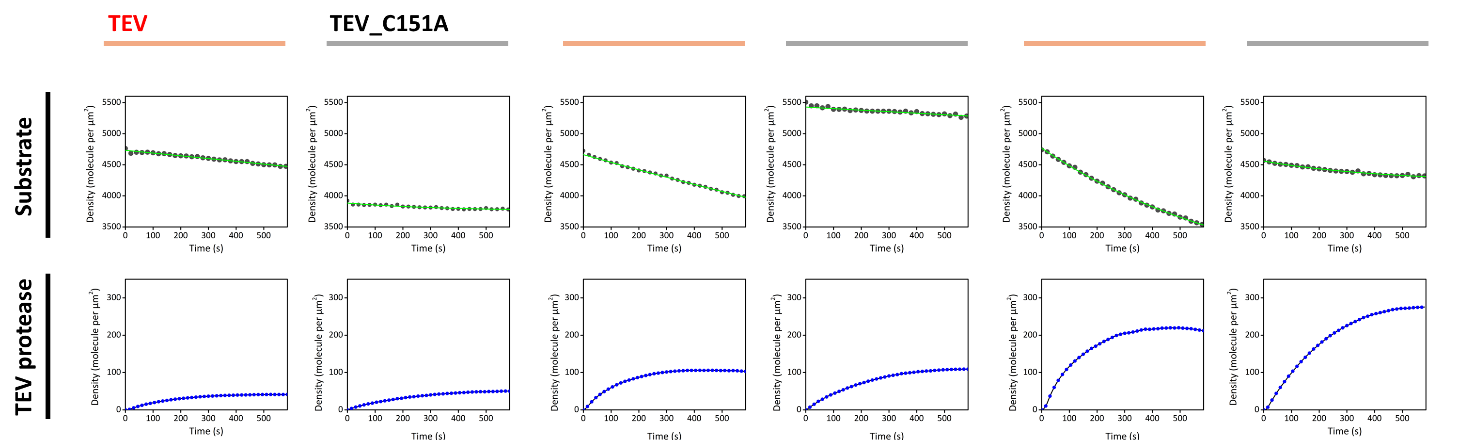

Figure S4. A control experiment to determine the extent of competition for the Ni-NTA binding site between TEV and GFP causing positive bias. The inactive mutant TEV^C151A^ served as a control to measure the decrease in GFP fluorescence on the membrane due to competitive binding. A comparison of active and inactive TEV shows a decrease in GFP fluorescence under three conditions with densities of 50, 100, and 200 TEV per µm². In all three conditions, recruitment of TEV^C151A^ causes a slight decrease in GFP fluorescence compared to active TEV, confirming that GFP dissociation from the membrane is primarily due to the cleavage reaction and not competition.

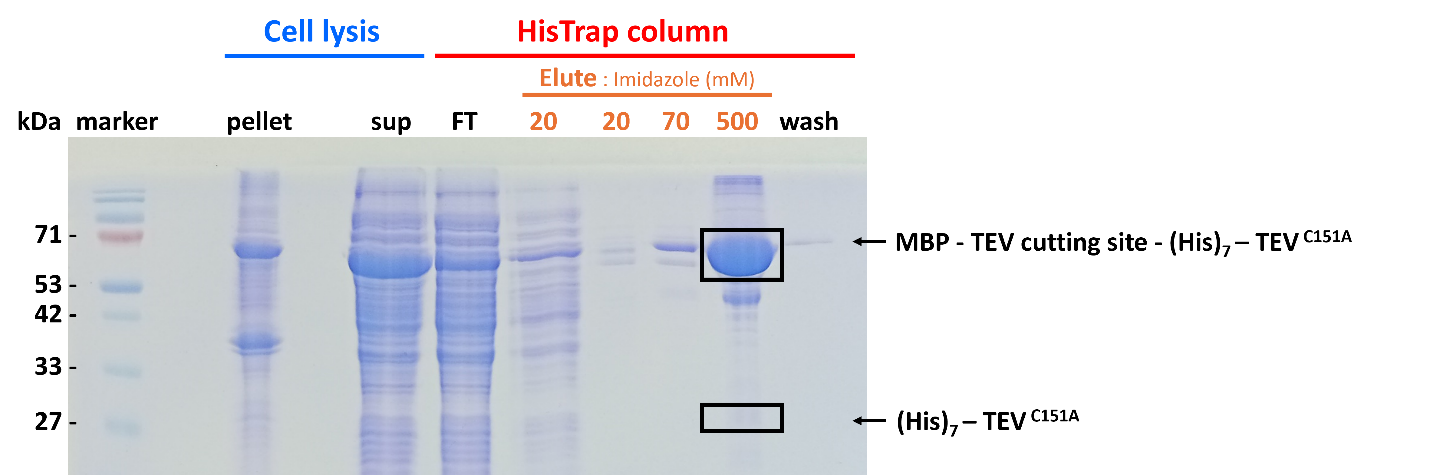

Figure S5. SDS-PAGE analysis of His7-TEV^C151A^ expression and purification. MBP and His7-TEV are fused by a TEV cleavage site. After expression, the MBP fusion tag is removed by self-cleavage and separated by a Ni-NTA column. The gel shows that the MBP tag remains fused with His7-TEV^C151A^ (71 kDa) after expression, confirming that TEV^C151A^ is inactive. No band appears at the 28 kDa position (TEV) in the SDS-PAGE.

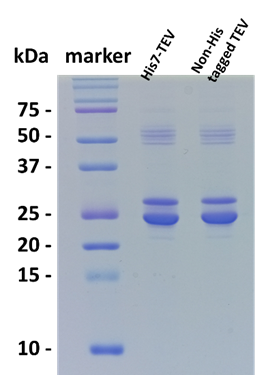

Figure S6. SDS-PAGE for comparison of His7-TEV and non-His-tagged TEV protease activities. TEV protease and substrate were mixed at a 1:50 (w/w) ratio, with 5.55 μg of TEV protease and 277 μg of substrate in a total volume of 40 μl. The reaction mixture was incubated at room temperature for 3 hours, and a 1 μl sample was taken for SDS-PAGE analysis. Lane 1: standard protein marker, Lane 2: product cleaved by His7-TEV, Lane 3: product cleaved by non-His-tagged TEV. This gel demonstrates that His7-TEV and non-His-tagged TEV exhibit comparable activities.

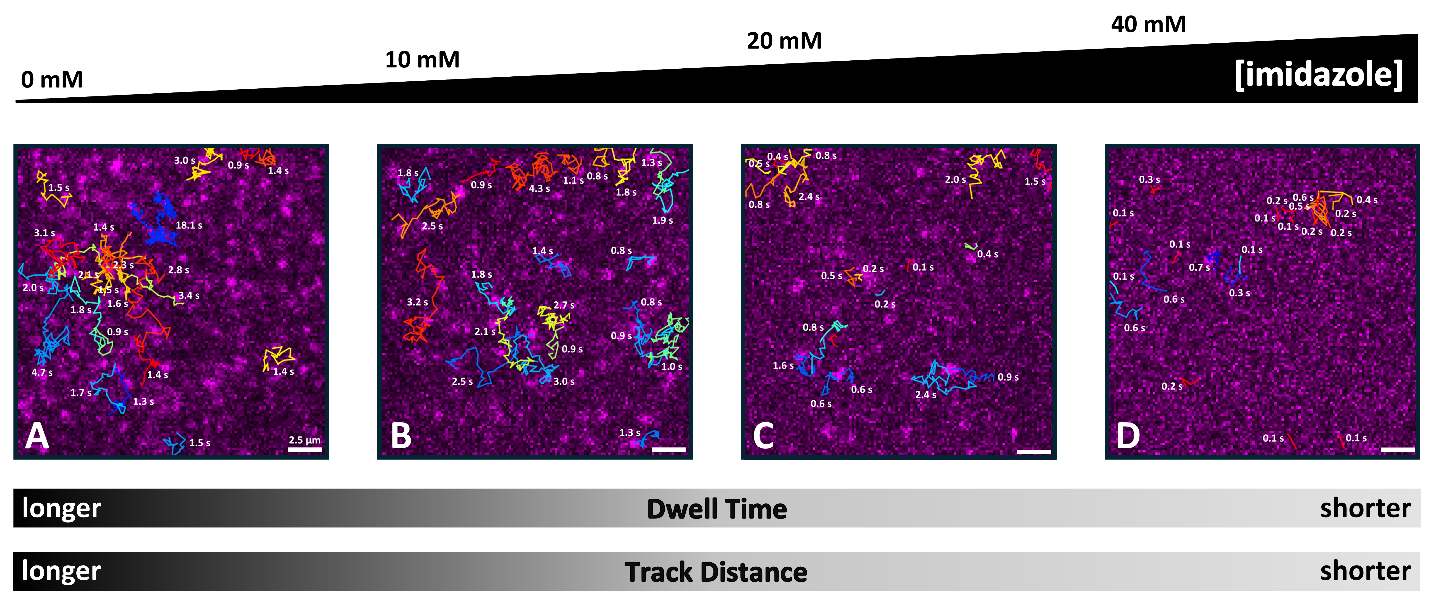

Figure S7. Analysis of membrane dwell time of His4-TEV by single-molecule tracking. TIRF imaging of fluorescently labeled (Alexa Fluor 647) TEV with (A) 0 mM, (B) 10 mM, (C) 20 mM, and (D) 40 mM imidazole was analyzed using single-molecule tracking. Partial single-molecule 2D diffusion trajectories of TEV are shown in the images (color-coded by track ID), with the corresponding dwell times labeled near each track. Single-molecule dwell time analysis shows a trend where higher imidazole concentrations result in shorter dwell times. (To prevent overcrowding in the image, only about 20 tracks filtered by track mean quality are shown.)
